# Microglia replacement effectively attenuates the disease progress of ALSP in the mouse model and human patient

**DOI:** 10.1101/2024.06.11.598496

**Authors:** Jingying Wu, Yafei Wang, Xiaoyu Li, Yuanyuan Cai, Pei Ouyang, Yang He, Mengyuan Zhang, Xinghua Luan, Yuxiao Jin, Taohui Liu, Yuxin Li, Xiaojun Huang, Ying Mao, Yanxia Rao, Li Cao, Bo Peng

## Abstract

Microglia play critical roles in the brain physiology and pathology. CSF1R is primarily expressed in microglia. The mono-allelic *CSF1R* mutation causes adult-onset leukoencephalopathy with axonal spheroids and pigmented glia (ALSP), a lethal neurological disease and no rational cure in clinical trials. There are no animal models mimicking human ALSP. In this study, we first developed mouse models based on human ALSP hotspot mutations. We then utilized microglia replacement by bone marrow transplantation (Mr BMT) to replace the *Csf1r*-deficient microglia in ALSP mice by *Csf1r*-normal donor cells. With pathogenic gene correction, Mr BMT efficiently attenuated the pathologies. Previously, an ALSP patient received traditional bone marrow transplantation (tBMT) due to a misdiagnosis of metachromatic leukodystrophy. The disease progress was halted for 15 years with unknown reasons. We demonstrated that tBMT in ALSP is equivalent to or close to Mr BMT, achieving efficient microglia replacement and therefore attenuating the ALSP progress in the mouse model. Next, we applied tBMT to replace *CSF1R*-deficient microglia in human patients. Our clinical results show that after microglia replacement, the ALSP course is effectively halted. Together, microglia replacement corrects the pathogenic gene and thus halts the disease progress in the mouse model and human patients. This study strongly demonstrates clinical potentials of microglia replacement in neurological disease treatments.

## Introduction

Microglia are important immune cells in the central nervous system (CNS). The dysfunction of microglia may cause or accelerate the CNS disorders. Colony-stimulating factor 1 receptor (CSF1R) is a tyrosine kinase^1^ that exclusively expresses in the brain myeloid cells, including the brain microglia (majority) and board-associated macrophages (BAMs) (minority). The survival of microglia relies on the CSF1R signaling^2,3^. The bi-allelic *CSF1R* mutation in human and mouse results in a congenital absence of microglia and is pediatrically lethal^2,4^. In contrast, the mono-allelic *CSF1R* mutation in human causes adult-onset leukoencephalopathy with axonal spheroids and pigmented glia (ALSP)^5^. This CSF1R-associated microgliopathy (CAMP) is a lethal neurological disease with microglial number reduction, myelin pathology (including hypermyelination and demyelination), brain parenchymal calcification and motor impairment^5–7^. There is no rational cure for ALSP. For historical reasons, ALSP is also known as CSF1R-related leukoencephalopathy (CRLE), hereditary diffuse leukoencephalopathy with axonal spheroids (HDLS), and pigmentary orthochromatic leukodystrophy (POLD)^8,9^.

ALSP is an overlooked disease and the mechanisms are poorly understood. One of the major reasons is the lack of adequate animal models mimicking human ALSP. To better study ALSP, we first generated CSF1R^WT/I792T^ and CSF1R^WT/E631K^ mice based on two hotspot mutations in human patients, *CSF1R*:NM_005211:c.2381T>C:p.I794T (the most common mutation worldwide^10–12^) and c.1897G>A:p.E633K (another common mutation in Europe and North America^11,12^). We demonstrated that these two models can mimic the human ASLP phenotypes with microglial number reduction, myelin pathology, brain parenchymal calcification and motor impairment.

In 2020, we first developed efficient strategies for microglia replacement^13^, which displays therapeutic potentials for neural disease treatment^14–19^. Since *CSF1R* gene mutation in microglia is the cause of ALSP, we therefore reasoned that replacing *CSF1R*-mutated microglia with *CSF1R*-normal microglia will be neuroprotective to ALSP patients. To test this hypothesis, we utilized microglia replacement by bone marrow transplantation (Mr BMT or mrBMT)^13,20,21^ to replace *Csf1r*-deficient microglia with *Csf1r*-normal microglia (or microglia-like cells) (refer to as replaced microglia or Mr BMT cells hereafter). We found that the gene correction by Mr BMT successfully enhances the microglia density with *Csf1r*-normal microglia. After microglia replacement, the ALSP symptoms were significantly alleviated including the myelin pathology, brain parenchymal calcification and motor impairment, demonstrating the therapeutic potential in pre-clinical studies.

In 2001, a patient was presumably diagnosed with adult-onset metachromatic leukodystrophy (MLD) and then received an allogeneic bone marrow cell (hematopoietic stem cell) transplantation from her unaffected brother, a standard procedure for MLD treatment^22^. After the traditional bone marrow transplantation (tBMT) for 15 years, the disease progress has been halted^7^. After the identification of ALSP-causing gene^5^, the patient was eventually diagnosed as ALSP with *CSF1R* mutation while her unaffected brother (the donor) is *CSF1R*-normal^7^. This is the first time showing tBMT can alleviate the ALSP development. After that, a few cases also show the neuroprotection of tBMT in ALSP^23^. However, the rationale is unknown. CSF1R signaling is a general pathway regulating the fitness and cell-cell competition of microglia^24^. *Csf1r*-deficient microglia are less-fitness cells that cannot compete the better-fitness *Csf1r*-normal microglia. In that scenario, tBMT in ALSP is equivalent to Mr BMT, which achieves an efficient replacement in the ALSP animal. Therefore, the therapeutic effects in these tBMT-treated ALSP patients are in fact from microglia replacement that corrects the pathogenic *CSF1R* mutations. To systematically evaluate the therapeutic effect in human, we performed microglia replacement in four ALSP patients by tBMT, replacing their *CSF1R*-deficient microglia with *CSF1R*-normal siblings. After microglia replacement, the disease progress has been halted for at least 24 months. It demonstrates the therapeutic effect of microglia replacement in treating human ALSP.

In summary, by combing pre-clinical and clinical studies, we demonstrated that microglia replacement can correct the disease-causing gene of ALSP and halt the disease progress in the animal model and human patients. It also indicates the therapeutic potentials of microglia replacement in treating other neurological disorders.

## Results

### Generation of CAMP model mimicking human ALSP symptoms

The adequate animal model is the prerequisite to study ALSP. To this end, we generated CSF1R^WT/I792T^ mice by CRISPR-based gene editing. The I792T mutation in mouse CSF1R corresponds to the I794T mutation in humans (*CSF1R*:NM_005211:c.2381T>C:p.I794T), which is the most common mutation worldwide^10–12^. CSF1R^WT/I792T^ mice showed significant microglial reductions from 6- to 16-month-old (Figure 1a-b and Figure S1). The brain calcification was observed in the CSF1R^WT/I792T^ mouse brain (Figure 1c). The CSF1R^WT/I792T^ mice displayed obvious motor impairments by the balance beam and rotarod tests (Figure 2a-c) and exhibited the ataxia-like behavior (Figure 2d). In addition to CSF1R^WT/I792T^, we also generated CSF1R^WT/E631K^ mice by CRISPR-based gene editing. The E631K mutation in mouse CSF1R corresponds to the E633K mutation in humans (*CSF1R*:NM_005211:c.1897G>A:p.E633K), which is another common mutation in Europe and North America^11,12^. We found that CSF1R^WT/E631K^ mice exhibited ALSP phenotypes with a microglial number reduction (Figure S2a-b and Figure S3) and brain calcification (Figure S2c). The motor impairment was observed in the balance beam and hindlimb clasping tests but not in the rotarod test (Figure S4). Collectively, by generating CSF1R mutant mice corresponding to human hotspot mutations, we built ALSP mouse models that mimic the human ALSP symptoms.

**Figure 1.**
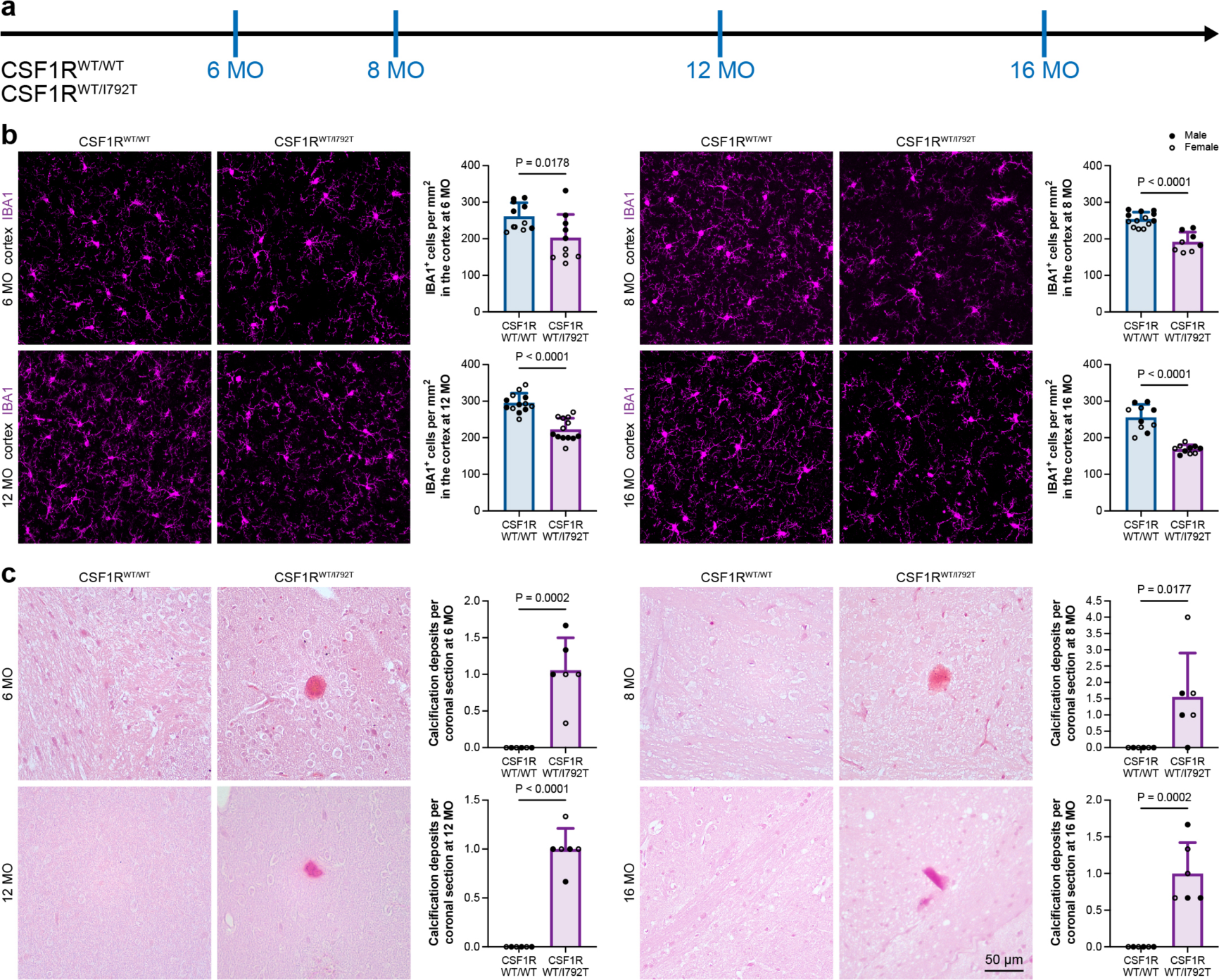
CSF1R^WT/I792T^ mice display pathological traits in the brain. **(a)** Scheme of the time point for examination. **(b)** CSF1R^WT/I792T^ mice exhibit a reduced microglial cell number in the cortex from 6- to 16-month-old. N = 11 (6-month-old), 13 (8-month-old), 13 (12-month-old) and 10 (16-month-old) mice for the CSF1R^WT/WT^ group, 10 (6-month-old), 8 (8-month-old), 13 (12-month-old) and 10 (16-month-old) mice for the CSF1R^WT/I792T^ group, respectively. The quantitative results of CSF1R^WT/WT^ mice are the same data in Figure 1b. **(c)** CSF1R^WT/I792T^ mouse brain exhibits calcification. N = 6 mice for each group. N = 6 mice for each group. Data are presented as mean ± SD. Male mice are indicated as solid dots and female mice are indicated as circled dots. Two-tailed independent t test.

**Figure S1.**
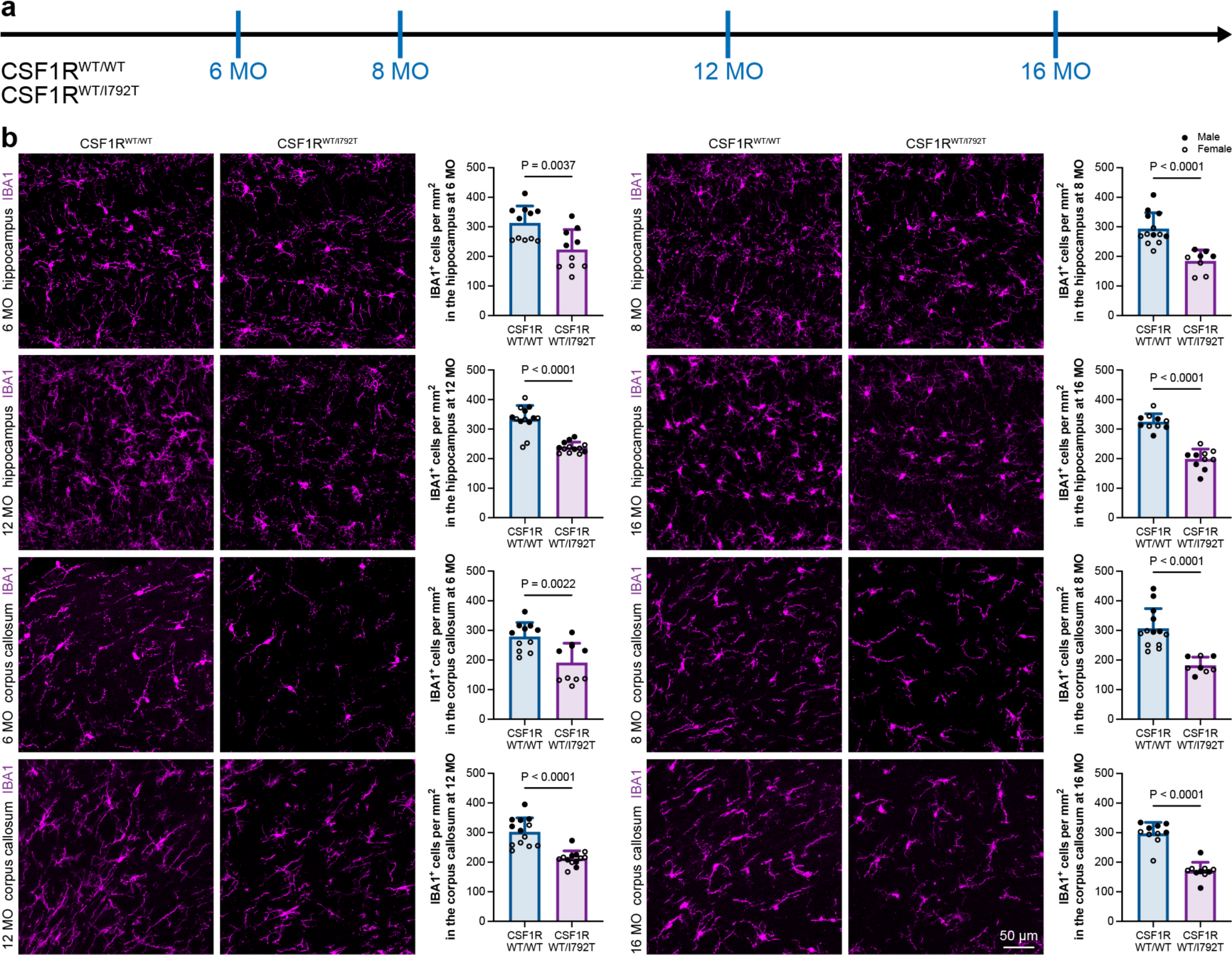
CSF1R^WT/I792T^ mice display microglial number reductions in the hippocampus and corpus callosum. **(a)** Scheme of the time point for examination. **(b)** CSF1R^WT/I792T^ mice exhibit a reduced microglial cell number in the hippocampus and corpus callosum from 6- to 16-month-old. N = 11 (6-month-old), 13 (8-month-old), 13 (12-month-old) and 10 (16-month-old) mice for the CSF1R^WT/WT^ group, 10 (6- month-old), 8 (8-month-old), 13 (12-month-old) and 10 (16-month-old) mice for the CSF1R^WT/I792T^ group, respectively. Data are presented as mean ± SD. Male mice are indicated as solid dots and female mice are indicated as circled dots. Two-tailed independent t test.

**Figure 2.**
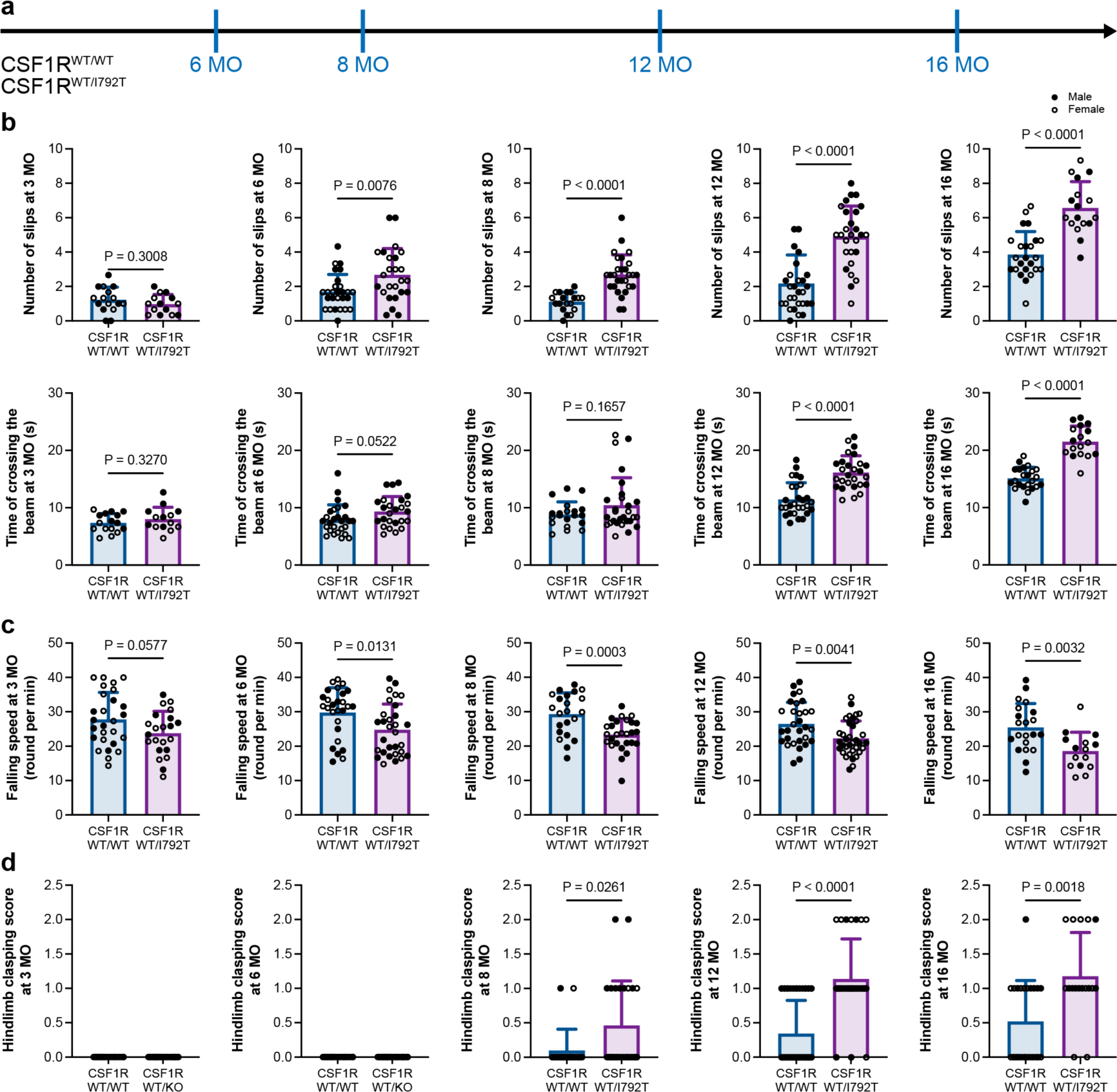
CSF1R^WT/I792T^ mice exhibit motor impairments. **(a)** Scheme of the time point for examination. **(b)** CSF1R^WT/I792T^ mice display significantly increased slip number from 6- to 16- month-old and increased crossing time from 12- to 16-month-old in the balance beam test. N = 16 (3-month-old), 27 (6-month-old), 19 (8-month-old, slip), 20 (8-month-old, crossing time), 28 (12-month-old), 24 (16-month-old, slip) and 23 (16-month-old, crossing time) mice for the CSF1R^WT/WT^ group, 14 (3-month-old), 25 (6-month-old), 28 (8-month-old), 27 (12-month-old) and 17 (16-month-old) mice for the CSF1R^WT/I792T^ group, respectively. **(c)** CSF1R^WT/I792T^ mice show reduced falling speed from 6- to 16-month-old in the rotarod test. N = 27 (3-month-old), 27 (6-month-old), 22 (8-month-old), 30 (12-month-old) and 21 (16-month-old) mice for the CSF1R^WT/WT^ group, 22 (3-month-old), 30 (6-month-old), 27 (8-month-old), 35 (12-month-old) and 15 (16-month-old) mice for the CSF1R^WT/I792T^ group, respectively. **(d)** CSF1R^WT/I792T^ mice show the ataxia-like behavior from 8- to 16-month-old. N = 15 (3-month-old), 25 (6-month-old), 20 (8-month-old), 32 (12-month-old) and 23 (16- month-old) mice for the CSF1R^WT/WT^ group, 16 (3-month-old), 23 (6-month-old), 26 (8-month-old), 29 (12-month-old) and 17 (16-month-old) mice for the CSF1R^WT/I792T^ group, respectively. Data are presented as mean ± SD. Male mice are indicated as solid dots and female mice are indicated as circled dots. Two-tailed independent t test.

**Figure S2.**
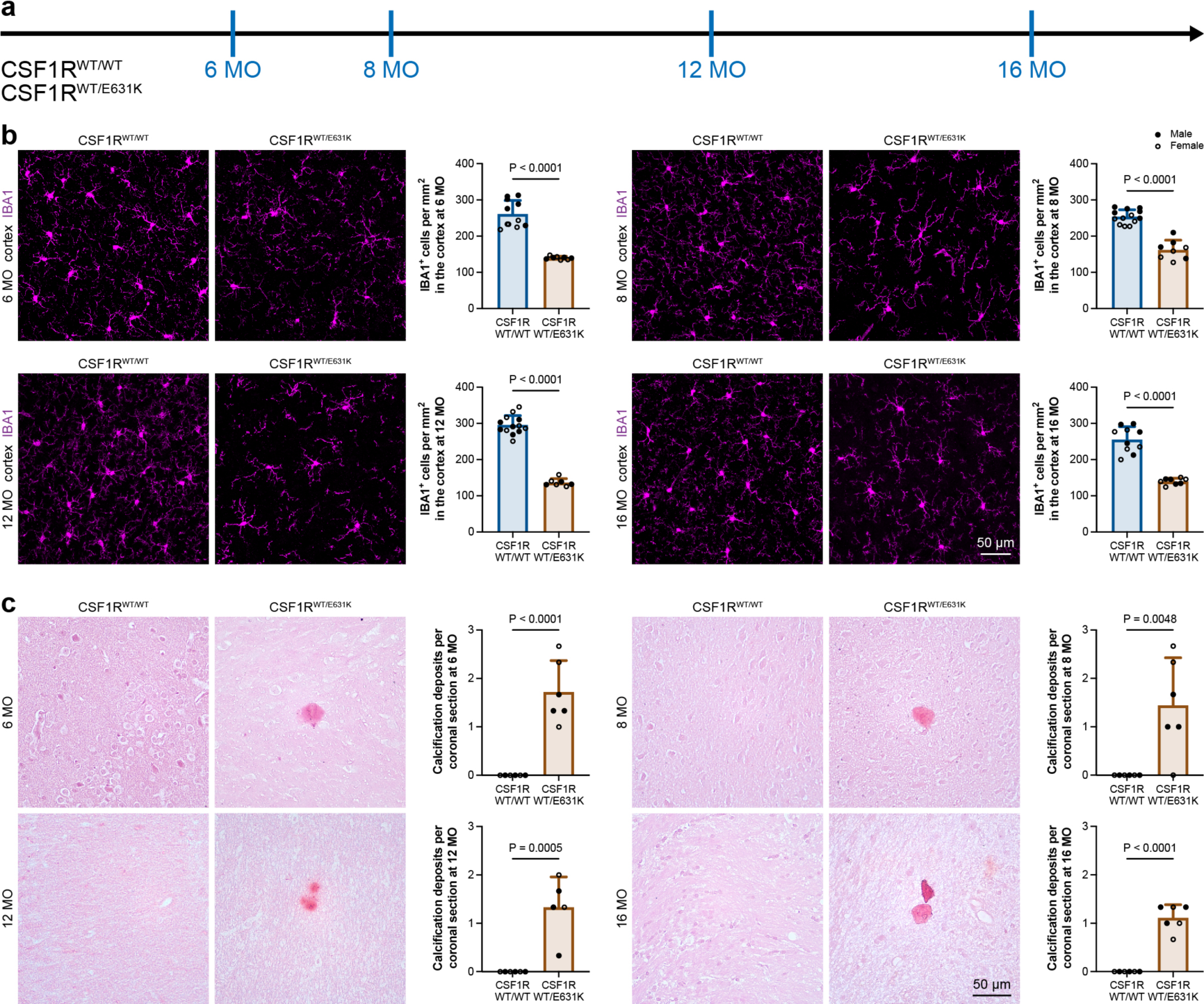
CSF1R^WT/E631K^ mice display pathological traits in the brain. **(a)** Scheme of the time point for examination. **(b)** CSF1R^WT/E631K^ mice exhibit a reduced microglial cell number in the cortex from 6- month-old to 16-month-old. N = 11 (6-month-old), 13 (8-month-old), 13 (12-month-old) and 10 (16-month-old) mice for the CSF1R^WT/WT^ group, 8 (6-month-old), 8 (8- month-old), 7 (12-month-old) and 8 (16-month-old) mice for the CSF1R^WT/E631K^ group, respectively. The quantitative results of CSF1R^WT/WT^ mice are the same data in Figure 1c. **(c)** CSF1R^WT/E631K^ mouse brain exhibits calcification. N = 6 mice for each group. Data are presented as mean ± SD. Male mice are indicated as solid dots and female mice are indicated as circled dots. Two-tailed independent t test. The quantitative results of CSF1R^WT/WT^ mice are the same data in Figure 1.

**Figure S3.**
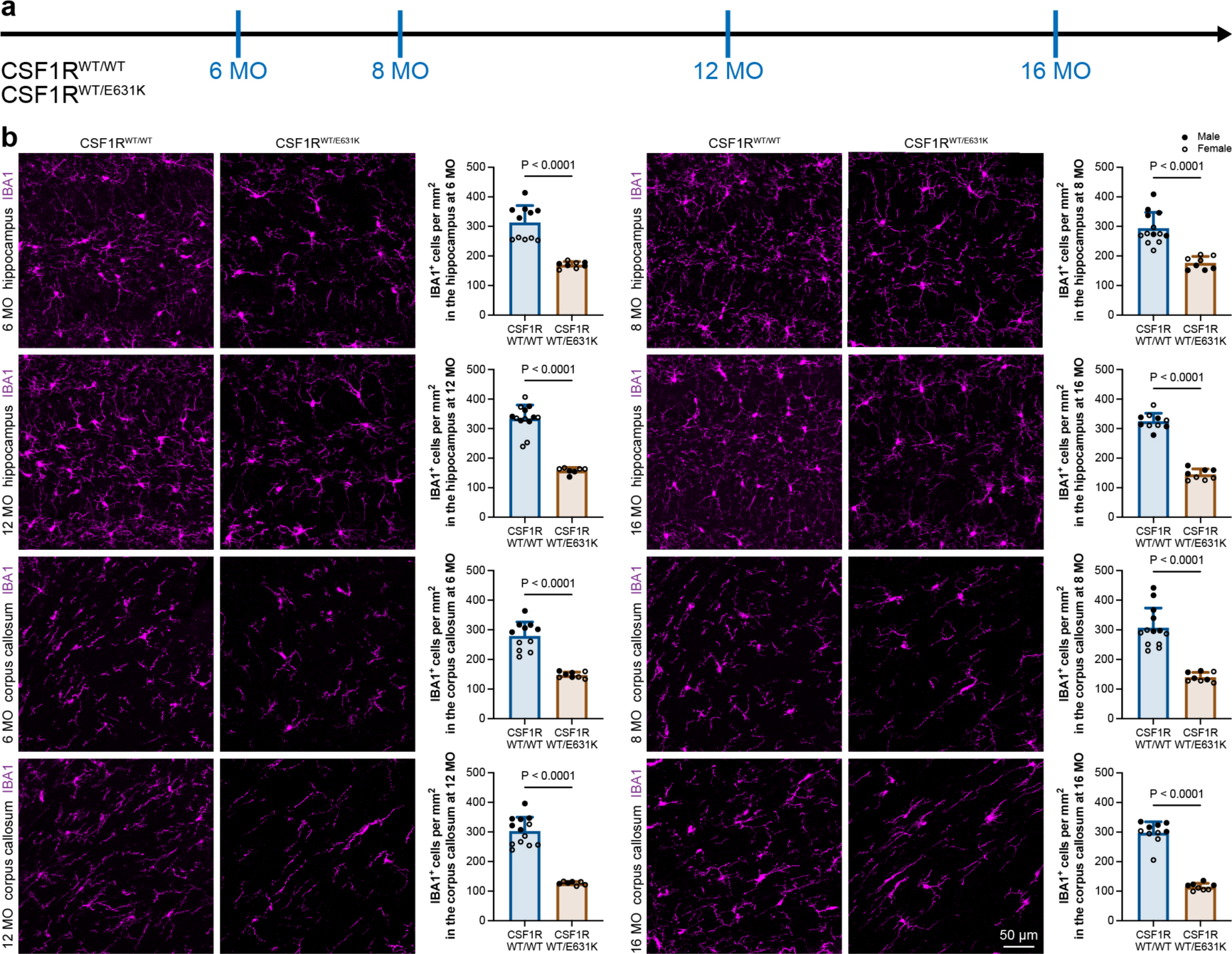
CSF1R^WT/E631K^ mice display microglial number reductions in the hippocampus and corpus callosum. **(a)** Scheme of the time point for examination. **(b)** CSF1R^WT/E631K^ mice exhibit a reduced microglial cell number in the hippocampus and corpus callosum from 6-month-old to 16-month-old. N = 11 (6-month-old), 13 (8- month-old), 13 (12-month-old) and 10 (16-month-old) mice for the CSF1R^WT/WT^ group, 8 (6-month-old), 8 (8-month-old), 7 (12-month-old) and 8 (16-month-old) mice for the CSF1R^WT/E631K^ group, respectively. The quantitative results of CSF1R^WT/WT^ mice are the re-analysis of same data in Figure S1b. Data are presented as mean ± SD. Male mice are indicated as solid dots and female mice are indicated as circled dots. Two-tailed independent t test.

**Figure S4.**
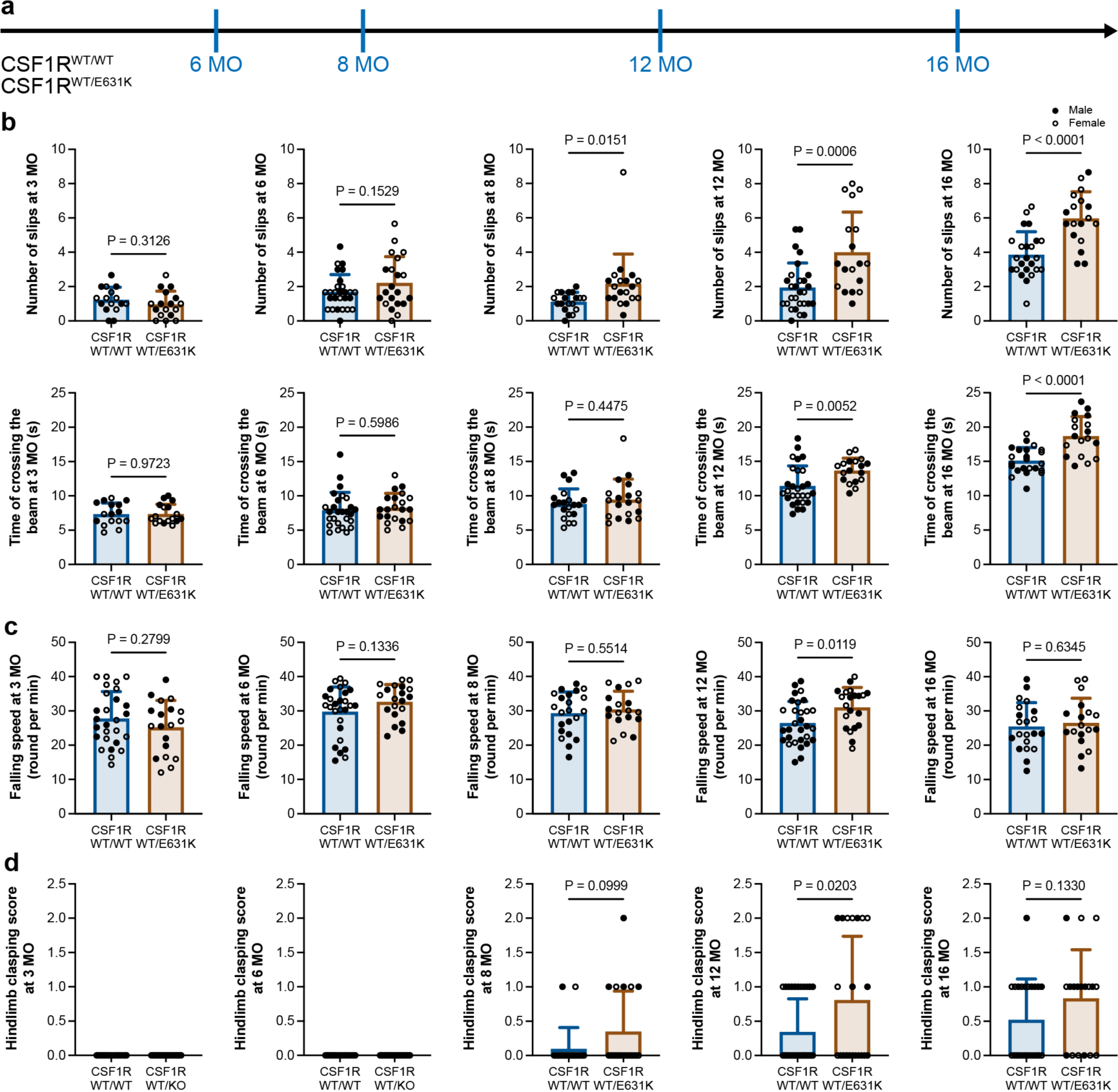
CSF1R^WT/E631K^ mice exhibit motor impairments. **(a)** Scheme of the time point for examination. **(b)** CSF1R^WT/E631K^ mice display significantly increased slip number from 8 to 16- month-old and increased crossing time from 12 to 16-month-old in the balance beam test. N = 16 (3-month-old), 27 (6-month-old), 19 (8-month-old, slip), 20 (8-month-old, crossing time), 28 (12-month-old), 24 (16-month-old, slip) and 23 (16-month-old, crossing time) mice for the CSF1R^WT/WT^ group, 17 (3-month-old), 21 (6-month-old), 19 (8-month-old), 18 (12-month-old) and 18 (16-month-old) mice for the CSF1R^WT/E631K^ group, respectively. **(c)** CSF1R^WT/E631K^ mice at 12-month-old show altered falling speed in the rotarod test. N = 27 (3-month-old), 26 (6-month-old), 22 (8-month-old), 30 (12-month-old) and 21 (16-month-old) mice for the CSF1R^WT/WT^ group, 19 (3-month-old), 20 (6-month-old), 18 (8-month-old), 21 (12-month-old) and 18 (16-month-old) mice for the CSF1R^WT/E631K^ group, respectively. **(d)** CSF1R^WT/E631K^ mice show the ataxia-like behavior at 12-month-old. N = 27 (3- month-old), 27 (6-month-old), 22 (8-month-old), 30 (12-month-old) and 21 (16-month-old) mice for the CSF1R^WT/WT^ group, 19 (3-month-old), 20 (6-month-old), 20 (8- month-old), 21 (12-month-old) and 18 (16-month-old) mice for the CSF1R^WT/E631K^ group, respectively. Data are presented as mean ± SD. Male mice are indicated as solid dots and female mice are indicated as circled dots. Two-tailed independent t test. The quantitative results of CSF1R^WT/WT^ mice are the same data in Figure 2.

### Mr BMT corrects the pathogenic gene and attenuates the disease progression in ALSP mice

We found that the gene correction by Mr BMT can attenuate the pathologies in the ALSP mouse brain (data will be shown soon). Next, we asked whether the pathological mitigation by Mr BMT can improve the motor performance of CSF1R^WT/I792T^ mice. To this end, we accessed the motor performance by the balance beam, rotarod and hindlimb clasping tests (Figure 3a). Mr BMT-treated CSF1R^WT/WT^ mice (normal replaced by normal) did not display better motor performances in the balance beam (Figure 3b, Video 1-2), rotarod (Figure 3c, Video 3) or hindlimb clasping (Figure 3d, Video 4-5). The results reveal that Mr BMT *per se* (without gene correction) does not enhance the motor performances. In contrast, CSF1R^WT/I792T^ mice received Mr BMT treatment (deficient replaced by normal) display better motor performances compared to naïve CSF1R^WT/I792T^ mice, including the less slip number in the balance beam from 6- to 16-month-old, shorter crossing time in the balance beam at 3-, 8-, 12- and 16-month-old (Figure 3b, Video 6-7), higher falling speed in the rotarod from 3- to 16-month-old (Figure 3c, Video 3), and significantly lower hindlimb clasping score at 12-month-old and a lower trend at 16-month-old (P = 0.0862) (Figure 3d, Video 8-9). Therefore, correction of the pathogenic gene by Mr BMT alleviates the motor impairments of ALSP mice.

**Figure 3.**
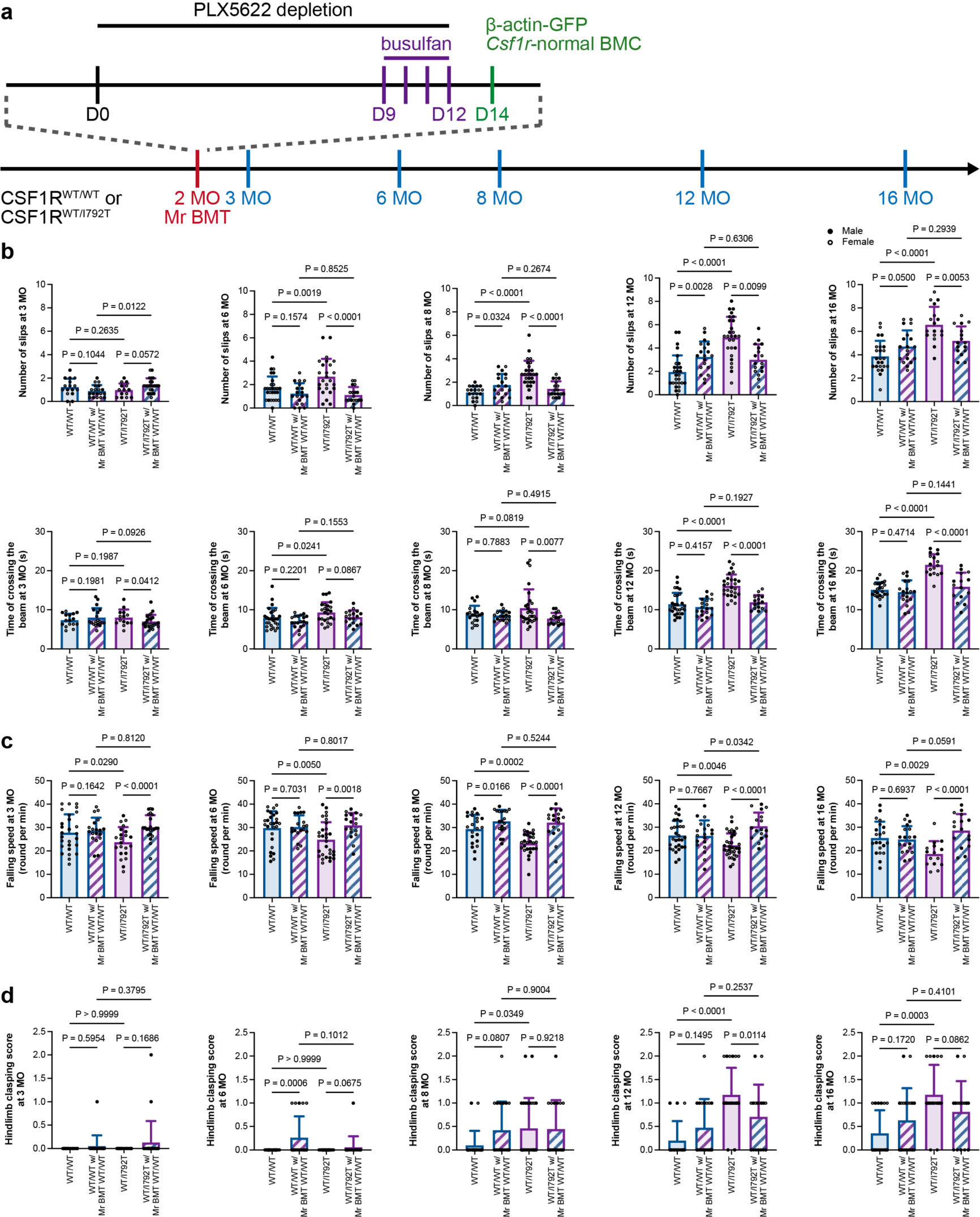
Mr BMT at 2-month-old does not enhance the motor performance in CSF1R^WT/WT^ mice while it significantly rescues the motor impairments in CSF1R^WT/I792T^ mice. **(a)** Scheme of the Mr BMT treatment and time points of behavior tests. **(b)** Mr BMT does not enhance the motor performance of CSF1R^WT/WT^ mice in the balance beam test. In contrast, CSF1R^WT/I792T^ mice received Mr BMT treatment display significantly less slip numbers from 6- to 16-month-old and shorter crossing time at 3-, 8-, 12- and 16-month-old. N = 16 (3-month-old), 27 (6-month-old), 19 (8-month-old, slip), 20 (8-month-old, crossing time), 28 (12-month-old), 24 (16-month-old, slip) and 23 (16-month-old, crossing time) mice for the naïve CSF1R^WT/WT^ group, 19 (3-month-old), 19 (6-month-old), 19 (8-month-old), 19 (12-month-old) and 19 (16-month-old) mice for the Mr BMT-treated CSF1R^WT/WT^ group, 22 (3-month-old), 30 (6-month-old), 27 (8-month-old), 35 (12-month-old) and 15 (16-month-old) mice for the naïve CSF1R^WT/I792T^ group, 23 (3-month-old), 18 (6-month-old), 18 (8-month-old), 17 (12- month-old) and 16 (16-month-old) mice for the Mr BMT-treated CSF1R^WT/I792T^ group, respectively. **(c)** Mr BMT does not enhance the motor performance of CSF1R^WT/WT^ mice in the rotarod test. In contrast, CSF1R^WT/I792T^ mice received Mr BMT treatment display significantly higher falling speed from 3- to 16-month-old. N = 27 (3-month-old), 27 (6-month-old), 22 (8-month-old), 30 (12-month-old) and 21 (16-month-old) mice for the naïve CSF1R^WT/WT^ group, 19 (3-month-old), 19 (6-month-old), 19 (8-month-old), 19 (12-month-old) and 19 (16-month-old) mice for the Mr BMT-treated CSF1R^WT/WT^ group, 22 (3-month-old), 30 (6-month-old), 27 (8-month-old), 35 (12-month-old) and 15 (16-month-old) mice for the naïve CSF1R^WT/I792T^ group, 23 (3-month-old), 18 (6- month-old), 18 (8-month-old), 17 (12-month-old) and 16 (16-month-old) mice for the Mr BMT-treated CSF1R^WT/I792T^ group, respectively. **(d)** Mr BMT does not enhance the motor performance of CSF1R^WT/WT^ mice in the hindlimb clasping test. In contrast, CSF1R^WT/I792T^ mice received Mr BMT treatment display a significantly lower hindlimb clasping score at 12-month-old and a lower trend at 16-month-old (P = 0.0862). N = 15 (3-month-old), 25 (6-month-old), 20 (8-month-old), 32 (12-month-old) and 23 (16-month-old) mice for the naïve CSF1R^WT/WT^ group, 19 (3-month-old), 19 (6-month-old), 19 (8-month-old), 19 (12-month-old) and 19 (16- month-old) mice for the tBMT-treated CSF1R^WT/WT^ group, 16 (3-month-old), 23 (6- month-old), 26 (8-month-old), 29 (12-month-old) and 17 (16-month-old) mice for the naïve CSF1R^WT/I792T^ group, 23 (3-month-old), 18 (6-month-old), 18 (8-month-old), 17 (12-month-old) and 16 (16-month-old) mice for the tBMT-treated CSF1R^WT/I792T^ group, respectively. Data are presented as mean ± SD. Male mice are indicated as solid dots and female mice are indicated as circled dots. Two-tailed independent t test. The quantitative results of naïve CSF1R^WT/WT^ and naïve CSF1R^WT/I792T^ mice are the same data in Figure 2.

Taken together, our results demonstrated that correcting the pathogenic gene by Mr BMT effectively rescues the brain pathology and motor impairments of ALSP mice.

### scRNA-seq results reveal that Mr BMT reshapes ALSP oligodendrocytes to a more normal-like phenotype

To further characterize the influence of Mr BMT to the ALSP mouse brain, we harvested brains from CSF1R^WT/I792T^, naïve CSF1R^WT/I792T^ and Mr BMT-treated CSF1R^WT/I792T^ (deficient replaced by normal) mice for single-cell RNA-sequencing (scRNA-seq) (Figure 4a). To prevent the *ex vivo* reactivation that may influence the microglial transcriptome, we included a cocktail of transcriptional and translational inhibitors during the cell preparation procedure as previous described^25,26^. Fourteen cell types were identified in mouse brains according to cell-type-specific gene profiles^26,27^, including several clusters of microglia/macrophage were identified from CSF1R^WT/I792T^, naïve CSF1R^WT/I792T^ and Mr BMT-treated CSF1R^WT/I792T^ mice (Figure 4b). We thus subset microglia/macrophage for further analysis (Figure 4b-c). Microglia/macrophage includes *Csf1r*-normal microglia, *Csf1r*-deficient microglia, *Gfp*^+^ *Csf1r*-normal Mr BMT cells and BAMs (endogenous *Gfp*^−^ BAM 1 and replaced *Gfp*^+^ BAM 2) (Figure 4c). As *Csf1r*-deficient microglia are the pathogenic cells in ALPS, we compared the transcriptome between naïve CSF1R^WT/I792T^ and CSF1R^WT/WT^ microglia. Among the differentially expressed genes (DEGs) (|log_2_FC| > 0.25, adjust P < 0.05) between these two populations, 582 biological processes were annotated by gene ontology (GO). Several biological processes were related to ALSP pathology, including axonogenesis, neuron death, axon ensheathment, regulation of myelination, and negative regulation of immune system process (Figure 4d and Table 1), suggesting the microglia-mediated etiology. In contrast, after Mr BMT treatment, 731 biological processes were enriched in the DEGs of Mr BMT-treated CSF1R^WT/I792T^ vs naïve CSF1R^WT/I792T^ microglia, including axonogenesis, gliogenesis, neuron death, regulation of neuron death, and regulation of myelination (Figure 4e and Table 2). The therapeutic effect of Mr BMT may mediated by these processes.

**Figure 4.**
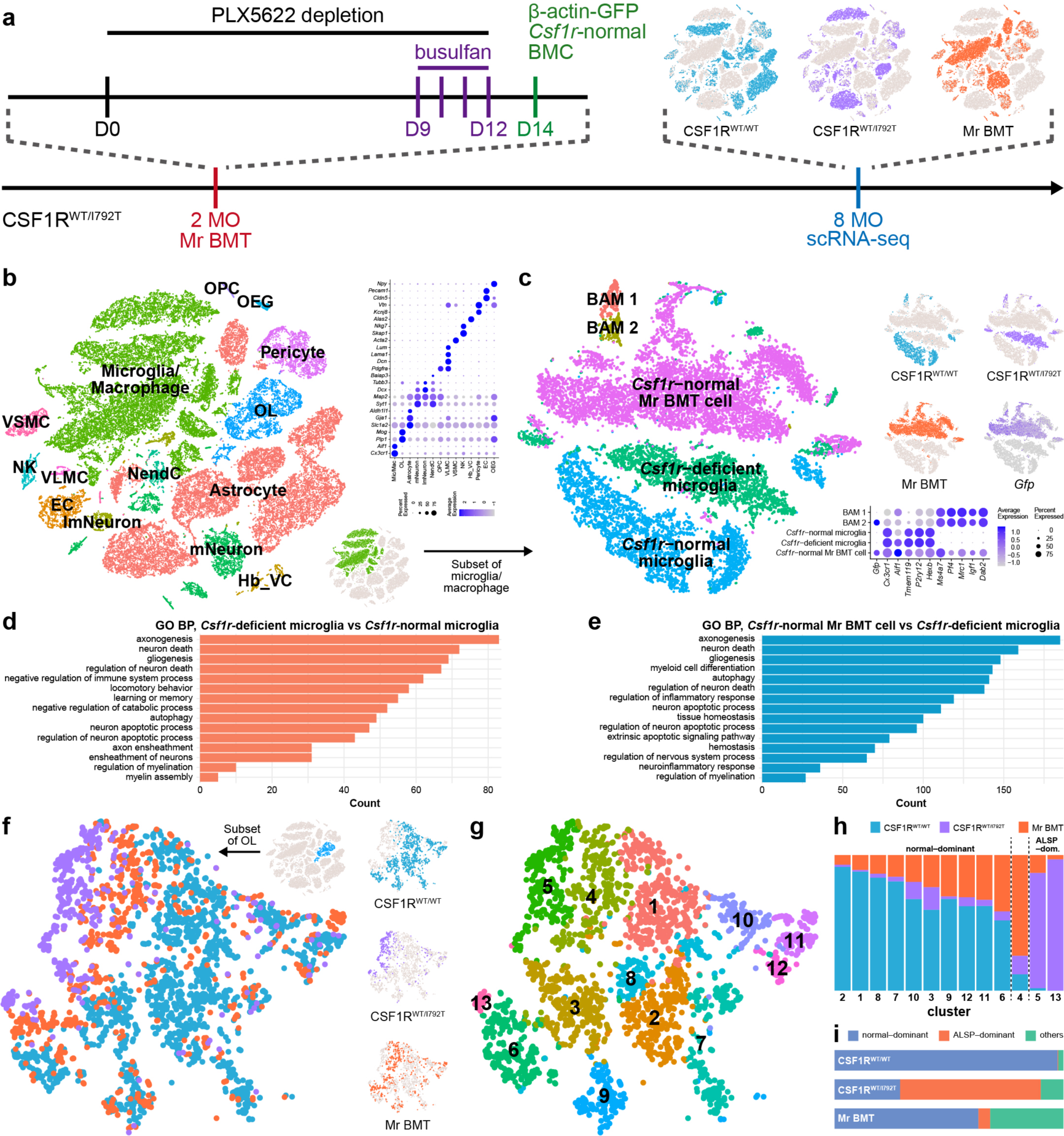
Mr BMT reshapes ALSP oligodendrocytes to a more normal-like phenotype. **(a)** Scheme of the Mr BMT treatment and time points of scRNA-seq. **(b)** Fourteen cell types were identified by scRNA-seq. **(c)** Microglia/macrophage subsets include *Csf1r*-normal microglia, *Csf1r*-deficient microglia, *Gfp*^+^ *Csf1r*-normal Mr BMT cells and BAMs (endogenous *Gfp*^−^ BAM 1 and replaced *Gfp*^+^ BAM 2). **(d-e)** Fifteen enriched GO biological processes of the *Csf1r*-deficient microglia vs *Csf1r*-normal microglia (d) and *Csf1r*-normal Mr BMT cell vs *Csf1r*-deficient microglia (e) DEGs. **(f-g)** The oligodendrocyte subsets are divided into 13 clusters. **(h)** The normal-dominant group includes clusters 2, 1, 8, 7, 10, 3, 9, 12, 11 and 6, whereas the ALSP-dominant group includes clusters 5 and 13. **(i)** Mr BMT treatment increases the oligodendrocyte percentage of ALSP mice in the normal-dominant clusters and decreases that in the ALSP-dominant clusters. BAM: boarder-associated macrophage; VLMC: vascular and leptomeningeal cell; Hb_VC: hemoglobin-expressing vascular cell; VSMC: vascular smooth muscle cell; EC: endothelial cell; OEG: olfactory ensheathing glia; mNeuron: mature neuron; ImmNeuron: immature neuron; NK: natural killer cell; NendC: neuroendocrine cell; OL: oligodendrocyte; OPC: oligodendrocyte precursor cell.

As myelin pathology is one of the hallmarks of ALSP and microglia are essential for myeline healthy^28,29^, we compared the influences of different microglia to the oligodendrocyte (OL) subset (Figure 4f). CSF1R^WT/I792T^ oligodendrocytes primarily located in the distant clusters from those of CSF1R^WT/WT^ mice (Figure 4f-g). We named these clusters as ALSP-dominant and normal-dominant clusters, respectively (Figure 4g-h). In the naïve CSF1R^WT/I792T^ mice, 61.47% of oligodendrocytes were in the ALSP-dominant clusters. In contrast, only 28.67% of naïve CSF1R^WT/I792T^ oligodendrocytes were in the normal-dominant clusters (Figure 4h-i). After Mr BMT treatment, only 5.17% of CSF1R^WT/I792T^ oligodendrocytes were in ALSP-dominant clusters whereas 62.86% of them resumed to the normal-dominant phenotype (Figure 4h-i). The results indicates that the gene correction by Mr BMT can reshape the ALSP oligodendrocytes to a more normal-like phenotype.

### tBMT in ALSP is equivalent to Mr BMT in replacement efficiency

The fitness and cell-cell competition of microglia is regulated by CSF1R^24^. CSF1R deficiency makes endogenous microglia in ALSP recipient less competitive against wild-type donor cells. As a result, tBMT in ALSP is equivalent to or close to Mr BMT, achieving efficient microglia replacement (data will be shown soon).

### Microglia replacement by tBMT corrects the pathogenic gene and mitigates the disease progress in ALSP mice

We found that microglia replacement by tBMT correcting the mutated *Csf1r* can attenuate the pathologies in the ALSP mouse brain (data will be shown soon). We then asked whether the tBMT-mediated mitigation of ALSP brain pathology can improve the motor performance in mice. We thus tested the motor performance through the balance beam, rotarod and hindlimb clasping tests (Figure 5a). tBMT-treated CSF1R^WT/WT^ mice (normal replaced by normal) did not display better motor performances in the balance beam (Figure 5b, Video 10), rotarod (except for at 3-month-old) (Figure 5c, Video 11) or hindlimb clasping (Figure 5d, Video 12). These results indicate that tBMT *per se* (non-gene correction) does not lead to a better motor performance. However, tBMT-treated CSF1R^WT/I792T^ mice (deficient replaced by normal) display improved motor performances compared to naïve CSF1R^WT/I792T^ mice, including a less slip number in the balance beam at 6-, 12- and 16-month-old, reduced crossing time in the balance beam from 12- to 16-month-old (Figure 5b, Video 13), higher falling speed in the rotarod from 3- to 16-month-old (Figure 5c, Video 11), and significantly lower hindlimb clasping score from 12- to 16-month-old (Figure 5d, Video 14). Thus, correction of the pathogenic gene via tBMT-achieved microglia replacement improves the motor performance of ALSP mice.

**Figure 5.**
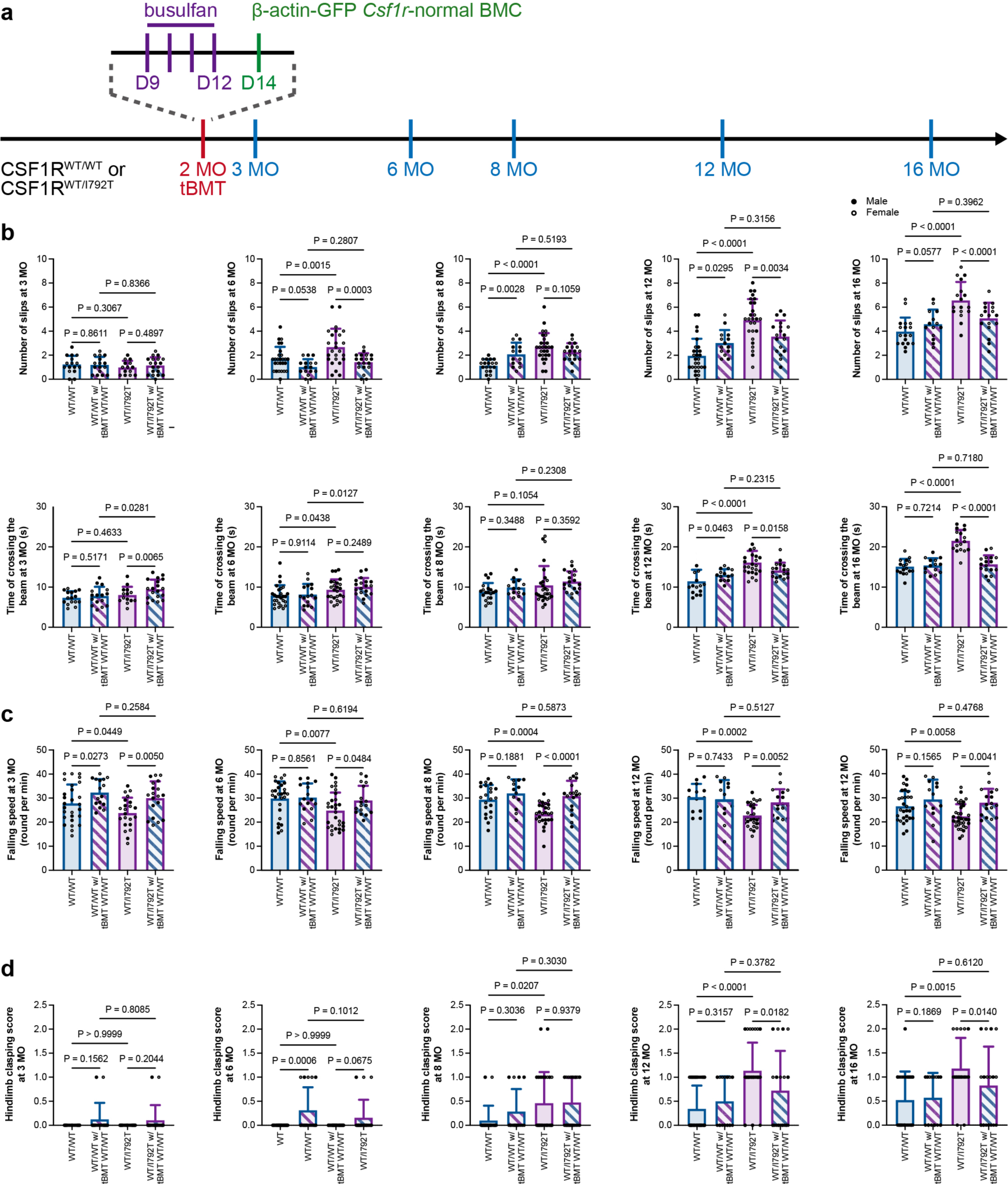
tBMT at 2-month-old does not enhance the motor performance in CSF1R^WT/WT^ mice while it significantly rescues the motor impairments in CSF1R^WT/I792T^ mice. **(a)** Scheme of the tBMT treatment and time points of behavior tests. **(b)** tBMT does not enhance the motor performance of CSF1R^WT/WT^ mice in the balance beam test. In contrast, CSF1R^WT/I792T^ mice received tBMT treatment display significantly less slip number at 6-, 12- and 16-month-old and shorter crossing time from 12- to 16-month-old. N = 16 (3-month-old), 27 (6-month-old), 19 (8-month-old, slip), 20 (8-month-old, crossing time), 28 (12-month-old), 24 (16-month-old, slip) and 23 (16-month-old, crossing time) mice for the naïve CSF1R^WT/WT^ group, 16 (3-month-old), 16 (6-month-old), 14 (8-month-old), 14 (12-month-old) and 14 (16-month-old) mice for the tBMT-treated CSF1R^WT/WT^ group, 16 (3-month-old), 23 (6-month-old), 26 (8-month-old), 29 (12-month-old) and 17 (16-month-old) mice for the naïve CSF1R^WT/I792T^ group, 19 (3-month-old), 19 (6-month-old), 19 (8-month-old), 18 (12- month-old) and 17 (16-month-old) mice for the tBMT-treated CSF1R^WT/I792T^ group, respectively. **(c)** tBMT does not enhance the motor performance of CSF1R^WT/WT^ mice in the rotarod test (except for at 3-month-old). In contrast, CSF1R^WT/I792T^ mice received Mr BMT treatment display a significantly higher falling speed from 3- to 16-month-old. N = 27 (3-month-old), 27 (6-month-old), 22 (8-month-old), 30 (12-month-old) and 21 (16- month-old) mice for the naïve CSF1R^WT/WT^ group, 16 (3-month-old), 16 (6-month-old), 13 (8-month-old), 13 (12-month-old) and 14 (16-month-old) mice for the tBMT-treated CSF1R^WT/WT^ group, 22 (3-month-old), 30 (6-month-old), 27 (8-month-old), 35 (12- month-old) and 15 (16-month-old) mice for the naïve CSF1R^WT/I792T^ group, 19 (3- month-old), 17 (6-month-old), 18 (8-month-old), 17 (12-month-old) and 17 (16-month-old) mice for the tBMT-treated CSF1R^WT/I792T^ group, respectively. **(d)** tBMT does not enhance the motor performance of CSF1R^WT/WT^ mice in the hindlimb clasping test. In contrast, CSF1R^WT/I792T^ mice received tBMT treatment display a significantly lower hindlimb clasping score from 12- to 16-month-old. N = 15 (3-month-old), 25 (6-month-old), 20 (8-month-old), 32 (12-month-old) and 23 (16- month-old) mice for the naïve CSF1R^WT/WT^ group, 16 (3-month-old), 16 (6-month-old), 14 (8-month-old), 14 (12-month-old) and 14 (16-month-old) mice for the tBMT-treated CSF1R^WT/WT^ group, 16 (3-month-old), 23 (6-month-old), 26 (8-month-old), 29 (12- month-old) and 17 (16-month-old) mice for the naïve CSF1R^WT/I792T^ group, 19 (3- month-old), 19 (6-month-old), 19 (8-month-old), 18 (12-month-old) and 17 (16-month-old) mice for the tBMT-treated CSF1R^WT/I792T^ group, respectively. Data are presented as mean ± SD. Male mice are indicated as solid dots and female mice are indicated as circled dots. Two-tailed independent t test. The quantitative results of naïve CSF1R^WT/WT^ and naïve CSF1R^WT/I792T^ mice are the same data in Figure 2.

Taken together, the *Csf1r*-deficiency results in a less-fitness in the cell-cell competition. In this scenario, tBMT achieves an efficient microglia replacement in *Csf1r*-deficienct ALSP mice, by which thus effectively attenuates the development of brain pathologies and improves the motor performance.

### Microglia replacement significantly halts the disease progress in ALSP human patients

To investigate whether microglia replacement can clinically halt or attenuate the progression of ALSP in human patients, we compared the magnetic resonance imaging (MRI) of 4 ALSP patients who received tBMT with HLA-matched *CSF1R*-normal cells (patients’ information see Supplementary Table 3) (Figure 6a) to the patients who did not receive tBMT. ALSP patients exhibited atrophy in the cortex and corpus callosum in T1 sequence, as well as periventricular white matter hyperintensities in T2-FLAIR and DWI (Figure 6b). MRI revealed the rapid disease progression in the tBMT-untreated ALSP patients with more severe atrophy in T1 and more hyperintensities in T2-FLAIR and DWI (Figure 6b), resulting in an increased Sundal MRI severity score (SSS)^30^ (Figure 6c). In contrast, microglia replacement by tBMT effectively halted the disease progress with cortical and corpus callosum atrophy no longer exacerbated in T1, and fewer hyperintensities in T2-FLAIR and DWI (Figure 6b). The SSS remains stable for the next 24 months after replacement (Figure 6c). Therefore, microglia replacement effectively halts the pathological progression in the ALSP patient’s brain.

**Figure 6.**
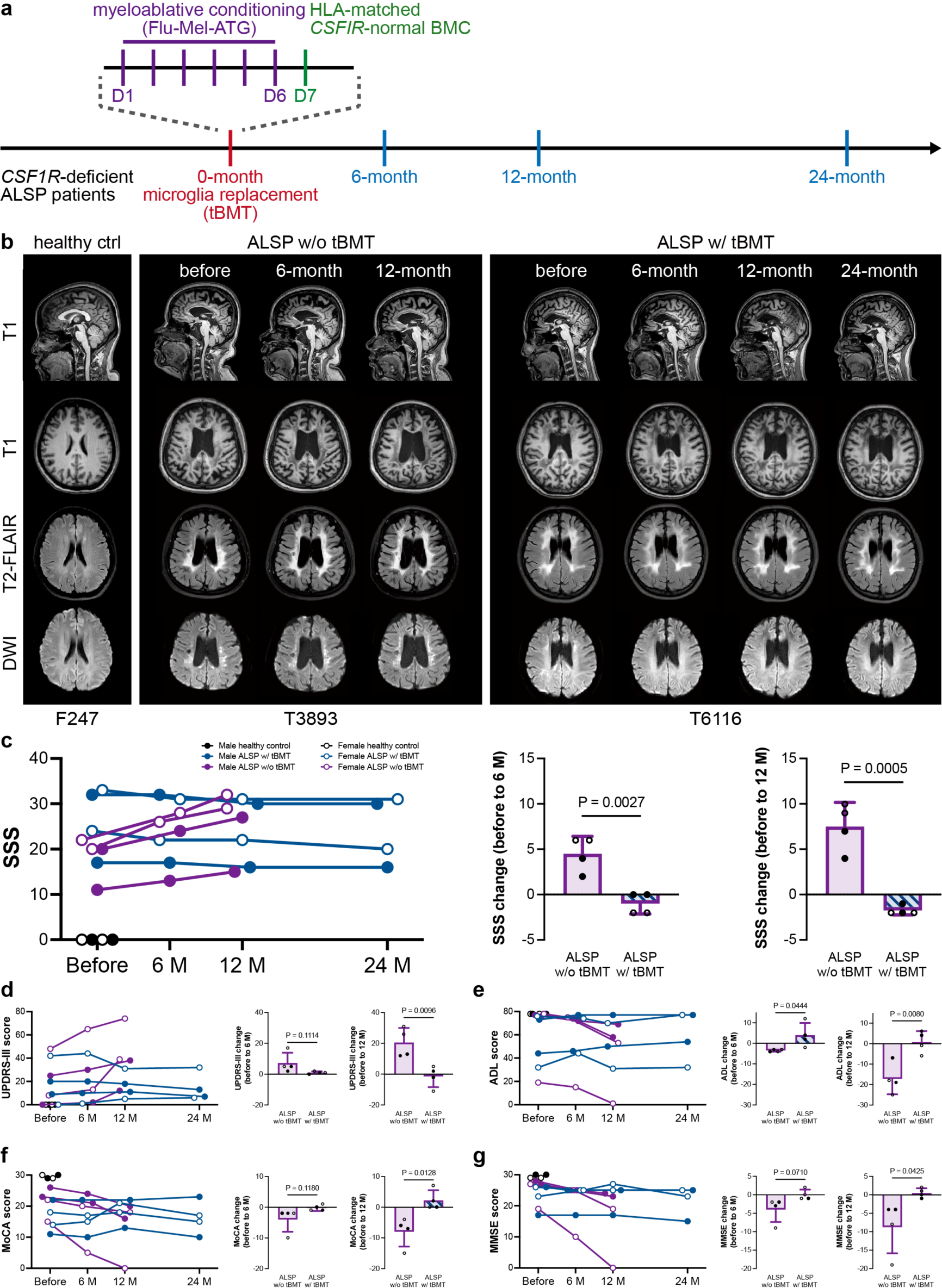
Microglia replacement by tBMT effectively halts the disease progress of ALSP patients. **(a)** Scheme of microglia replacement therapy by tBMT in human patients and following examinations. **(b-c)** MRI results reveal that microglia replacement effectively halts the pathological progress in human patients. **(d-g)** Microglia replacement effectively halts functional impairments of ALSP patients. Data in the histogram are presented as mean ± SD. Male patients are indicated as solid dots and female patients are indicated as circled dots. Two-tailed independent t test.

Next, we utilized unified Parkinson’s disease rating scale part-III (UPDRS-III) (a clinician-scored monitored motor evaluation, by which a higher score indicates a more severe motor impairment^31^) and activity of daily living (ADL) (the self-care tasks including bathing and showering, personal hygiene and grooming, dressing and toilet hygiene, functional mobility and self-feeding, by which a lower score indicates a worse performance in daily living activities^32^) to further evaluate the functional improvement in motor ability. Compare to the healthy controls, the ALSP patients exhibited higher UPDRS-III and lower ADL scores (Figure 6c-d). The motor impairments of the patients exacerbated rapidly in the following 12 months (Figure 6d-e). In contrast, microglia replacement by tBMT largely halted the progress of motor dysfunctions revealed by both of UPDRS-III (an improvement trend at 6-month post replacement, P = 0.1114, and significantly improved at 12-month post replacement) (Figure 6d) and ADL (significantly improved at 6- to 12-month post replacement) (Figure 6e). Twenty-four months after the tBMT, the motor function of ALSP patients remained stable and even showed a reverse trend (Figure 6d-e). ALSP patients also manifest with cognitive declines^6^. We thus used mini-mental state examination (MMSE) (a 30-point questionnaire measuring cognitive impairment, by which a lower score indicates a more severe cognitive decline^33,34^) and Montreal cognitive assessment (MoCA) (an evaluation including short-term memory, visuospatial abilities, multiple aspects of executive function, attention, concentration, working memory, language, abstract reasoning and orientation to time and place, by which a lower score indicates a more severe cognitive decline^35,36^) to evaluate the cognitive function. In contrast to the tBMT-nontreated patients, the tBMT-treated patients displayed a halted motor performance and an improvement trend compared to the ALSP patients who did not receive tBMT at 6-month after replacement in both MMSE (P = 0.1180) (Figure 6f) and MoCA (P = 0.0710) (Figure 6g). Twelve-month after replacement, the progress of cognitive impairment was significantly halted and displayed a significant improvement compared to the non-replaced patients in both (MMSE) and MoCA (Figure 6f-g). The cognitive function of tBMT-treated patients remained stable 24 months after microglia replacement (Figure 6f-g). Together, our results demonstrated that microglia replacement by tBMT effectively halted the development of functional impairment in ALSP patients.

## Supplementary

**Supplementary Table 1** Biological processes enriched in the DEGs of CSF1R^WT/WT^ vs naïve CSF1R^WT/I792T^ microglia.

**Supplementary Table 2** Biological processes enriched in the DEGs of Mr BMT-treated CSF1R^WT/I792T^ vs naïve CSF1R^WT/I792T^ microglia.

**Supplementary Table 3** Baseline characteristics of ALSP patients with or without tBMT and healthy controls.

## Videos

**Video 1** Naïve CSF1R^WT/WT^ mouse in the balance beam test.

**Video 2** Mr BMT-treated CSF1R^WT/WT^ mouse in the balance beam test.

**Video 3** Naïve CSF1R^WT/WT^, Mr BMT-treated CSF1R^WT/WT^, naïve CSF1R^WT/I792T^ and Mr BMT-treated CSF1R^WT/I792T^ mice in the rotarod test.

**Video 4** Naïve CSF1R^WT/WT^ mouse in the hindlimb clasping test.

**Video 5** Mr BMT-treated CSF1R^WT/WT^ mouse in the hindlimb clasping test.

**Video 6** Naïve CSF1R^WT/I792T^ mouse in the balance beam test.

**Video 7** Mr BMT-treated CSF1R^WT/I792T^ mouse in the balance beam test.

**Video 8** Naïve CSF1R^WT/I792T^ mouse in the hindlimb clasping test.

**Video 9** Mr BMT-treated CSF1R^WT/I792T^ mouse in the hindlimb clasping test.

**Video 10** tBMT-treated CSF1R^WT/WT^ mouse in the balance beam test.

**Video 11** Naïve CSF1R^WT/WT^, tBMT-treated CSF1R^WT/WT^, naïve CSF1R^WT/I792T^ and tBMT-treated CSF1R^WT/I792T^ mice in the rotarod test.

**Video 12** tBMT-treated CSF1R^WT/WT^ mouse in the hindlimb clasping test.

**Video 13** tBMT-treated CSF1R^WT/I792T^ mouse in the balance beam test.

**Video 14** tBMT-treated CSF1R^WT/I792T^ mouse in the hindlimb clasping test.

## Discussion

### Microglia replacement corrects the pathogenic gene and halts the disease progress of ALSP in mice and human patients

In 2020, we first developed three efficient strategies for microglia replacement, including Mr BMT^13,20^, microglia replacement by peripheral blood (Mr PB or mrPB)^13,37^ and microglia replacement by microglia transplantation (Mr MT or mrMT)^13,38^. We proposed potential applications in the neurological disease treatment including attenuating the progress of Alzheimer’s disease (AD) patients with the TREM2^R47H^ mutation^13^. Each strategy has its own advantages and can fit into different clinical scenarios^13–15^.

Evidences from mouse models strongly demonstrated the therapeutic potentials of microglia replacement in Alzheimer’s disease (AD), multiple sclerosis (MS) and a progressive neurodegeneration model^16–19^. These diseases, however, are not microgliopathy that with mutations in microglia specific and pathogenic causing genes. Thus, the proof from direct and rational treatment to microgliopathy is still absent. In this study, we established mouse models of CSF1R-associated microgliopathy, the major form of ALSP. We demonstrated that the correction of the pathogenic *Csf1r* mutation by microglia replacement is able to effectively mitigate the ALSP pathology and functional impairment in the mouse model. More importantly, based on the data of our clinical trials, we further demonstrated that microglia replacement can effectively halt the ALSP progress of human patients. This is a clinical proof of neurological disease treatment by microglia replacement. Microglia replacement therefore display great potentials in treating other neurological diseases, including microglia-causing diseases (microgliopathy) and microglia-accelerating diseases (e.g., AD with accompanying TREM2 mutation) (see Discussion in Xu et al., 2020, Cell Reports^13^).

Interestingly, microglia replacement also displayed therapeutic effects in an AD model without mutations of any microglia-expressing genes^18^. The rationale why the non-gene-correction microglia replacement protects AD brain is largely unknown. The replaced cells by Mr BMT and Mr PB are not identical to resident microglia. Compared to the yolk sac-derived *P2ry12*^+^ *Tmem119*^+^ *Siglech*^+^ *Mf4*^−^ *Ms4a7*^−^ microglia, the bone marrow cell derived-Mr BMT cells and blood cell-derived Mr PB cells are *P2ry12*^low^ *Tmem119*^low^ *Siglech*^low^ *Mf4*^+^ *Ms4a7*^+13^. In this study, the expression of microglia-specific and macrophage-specific genes of Mr BMT cells are between those of resident microglia and BAMs (Figure 4c). The unique phenotype might provide some neuroprotective effects. In addition, microglial reactivation and neurodegeneration are mutually enhanced. The inflamed microglia are removed in the first step of microglia replacement. The removal of microglia *per se* also can ameliorate the disease pathology. It is not only in the case of neuroprotective non-gene-correction microglia replacement, but may also in the case of neuroprotective microglia repopulation^39^.

### The cell-cell competition of microglia

Microglia display a cell-cell competition and do not “invade” the territory of their neighbors. In that way, microglia tile the brain and maintain a stable number in homeostasis. To achieve the efficient microglia, it is necessary to create a microglia-free niche and suppress the microglia proliferation in the recipient subjects^13^. The microglia-free niche can be achieved by CSF1R inhibition^3,13,40,41^ whereas suppression of microglia proliferation can be achieved by busulfan of irradiation^13,21,42–44^. This is echoed by a parabiosis study that allogenic myeloid cells differentiated into microglia and replaced the recipient’s microglia through a long-term ablation of CX3CR1-CreER::CSF1R^fl/fl^ microglia, by which simultaneously generated a microglia-free niche and inhibited the endogenous microglia without influencing donor cells^45^. Due to the lack of microglia-free niche in the physiological condition, tBMT is not capable of achieving an efficient microglia replacement^13^. However, the CSF1R-deficient ALSP is a partially microglia-free condition with reduction of microglial cell number and less cell-cell competitive^24^. In addition, busulfan or irradiation during the tBMT can further reduce the microglial number^42–44^. Therefore, ALSP is a natural microglia-free condition upon tBMT. Indeed, our results demonstrated that the replacement efficiency of tBMT in ALSP is equivalent or close to Mr BMT. It explains why the pathology can be halted in the misdiagnosed patient who received tBMT^7^. Furthermore, it also indicates that no extra CSF1R inhibition is required in the treatment of ALSP patients (that is tBMT).

On the other hand, for the other diseases that the microglial cell-cell competition is not impaired, microglia depletion is required for replacement. The safety of short- and long-term CSF1R inhibition has been well characterized in animals^3,13,40,46,47^ and human patients^48–51^. The microglial depletion by CSF1R inhibition has been granted US FDA approval^52,53^. Therefore, microglial replacement meets the safety requirements for clinical therapy. Even for some disease conditions, microglial depletion *per se* is neuroprotective^54–61^. Taken together, microglia replacement is a clinically feasible strategy showing great promising in other neurological disease treatment.

## Methods

### Ethics statements

All mouse experiments were conducted in accordance with the guidelines of the Institutional Animal Care and Use Committee of Fudan University, Animal Welfare Ethics Committee at Shanghai Sixth People’s Hospital Affiliated to Shanghai Jiao Tong University School of Medicine. This study was approved by the Institutional Animal Care and Use Committee of Fudan University (202001002Z and 2022JS-ITBR-023) and Animal Welfare Ethics Committee at Shanghai Sixth People’s Hospital Affiliated to Shanghai Jiao Tong University School of Medicine (2023-0126).

The clinical trial was approved by the Ethics Committee of Shanghai Sixth People’s Hospital (Approval No.: 2021-021, 2021-219), and registered in the China Clinical Trial Center (Registration No.: ChiCTR2100044666, ChiCTR2100050834).

### Animals

C57BL/6J mice were purchased from SPF (Beijing) Vital River Laboratory Animal Technology. β-actin-GFP mice (C57BL/6-Tg (CAG-EGFP) 131Osb/LeySopJ, Stock# 006567)^62^ and CSF1R^fl/fl^ (also known as CSF1R^tmdex5jwp^) mice (B6.Cg-*Csf1r^tm^*^1*.2Jwp*^/J, Stock# 021212)^63^ were purchased from The Jackson Laboratory. B6-CAG-Cre mice (B6/JGpt-Tg(CAG-Cre)/Gpt, Stock# T004055) were purchased from GemPharmatech China. CSF1R^WT/KO^ mice generated by crossing CSF1R^fl/fl^ mice with B6-CAG-Cre mice. All mice were housed in the Animal Facility at the Department of Laboratory Animal Science at Fudan University and Laboratory Animal Center at Shanghai Sixth People’s Hospital Affiliated to Shanghai Jiao Tong University School of Medicine under a 12-hour light/dark cycle with food and water were given *ad libitum*.

CSF1R^+/I792T^ (p.Ile792Thr, c.T2375C, ATT>ACT) mice were generated by Cyagen Biosciences China through CRISPR/Cas9-based extreme genome editing according to the authors’ design as a “fee-for-service”. The sgRNA was designed in the exon18, the targeting vector and donor oligo with targeting sequence, flanked by 123 bp homologous sequence combined on 5’ homologous arm and 3’ homologous arm. CSF1R^+/E631K^ (p.Glu631Lys, c.G1891A, GAG>AAG) mice were generated by Beijing Biocytogen China through CRISPR/Cas9-based extreme genome editing (EGE) according to the authors’ design as a “fee-for-service”. The sgRNA was designed in the intron 14, and the 5’ homologous arm and 3’ homologous arm for homologous recombination were approximately 1.3 kb and 1.1 kb, respectively.

Both male and female mice were utilized in this study. The sex of each mouse was indicated in the figure (male mice were indicated as solid dots and female mice were indicated as circled dots).

### Chemicals and reagents

Actinomycin D (HY-17559) was purchased from MedChemExpress. Anisomycin (HY-18982) was purchased from MedChemExpress. Busulfan (B2635) was purchased from Sigma-Aldrich. The normal AIN-76A diet (control diet, CD) was formulated by SYSE Bio (Cat#: PD1001, Lot#: 23020802). Ketamine (H20193336) was purchased from Shanghai Pharmaceutical. PLX5622 was formulated into the AIN-76A diet (1.2 g PLX5622 per kilogram of diet by SYSE Bio Cat#: D20010801, Lot#: 23021005). Triptolide (HY-32735) was purchased from MedChemExpress. Xylazine hydrochloride (Cat#: X1251) was purchased from Sigma-Aldrich.

### Drug administration

To pharmacologically eliminate myeloid cells, the mice were administered a PLX5622 (MCE)-formulated AIN-76A diet (1.2g PLX5622 per kilogram of diet formulated by SYSE Bio) *ad libitum* for 14 days. For microglia replacement, the mice were administrated 25 mg/kg busulfan by intraperitoneal injection for 4 days.

### Brain tissue preparation

Mice were deeply anesthetized with isoflurane. For histological experiments, animals were sequentially perfused with 0.01 M PBS and 4% paraformaldehyde (PFA) (Biosharp, Cat#: BL539A) in 0.01 M PBS. Brains were then carefully harvested and postfixed in 4% PFA in 0.01 MPBS at 4 °C overnight.

### Cryosection preparation

Brains and peripheral organs were dehydrated in 30% sucrose in 0.01 M PBS at 4 °C for 3-5 days. Samples were then embedded in optimal cutting temperature compound (OCT, SAKURA, Cat#: 4583), and frozen and stored at -80 °C before sectioning. Tissue with regions of interest was cut by a cryostat (Leica, CM1950) at a thickness of 30 μm.

### Immunohistochemistry and image acquisition

Brain and peripheral organ sections were rinsed with 0.01 M PBS 3 times for 10 to 15 min, and then blocked for 2 hours with 4% normal donkey serum (NDS, Jackson, Cat#: 017-000-121) in 0.01 M PBS containing 0.3% Triton X-100 (Aladdin, Cat#: T109026) (PBST) at room temperature (RT), followed by incubation with primary antibodies with 1% NDS in PBST at 4 °C overnight. After rinsing with PBST for 3 times, the samples were incubated with fluorescent dye-conjugated secondary antibodies at RT for 2 hours with 1% NDS in PBST with 4’,6-diamidino-2-phenylindole (DAPI, 1:1000, Sigma-Aldrich, D9542). Afterward, the samples were rinsed three times, and then mounted with anti-fade mounting medium (SouthernBiotech, Cat#: 0100-01).

Primary antibodies included rabbit anti-IBA1 (1:500, Wako, Cat#: 019-19741, Lot: LEP3218), goat anti-IBA1 (1:500, Abcam, Cat#: ab5076, Lot: GR3403958-1), rabbit anti-GFP (1:500, Invitrogen, Cat#: A11122, Lot: 2273763), goat anti-GFP (1:500, Abcam, Cat#ab6673, Lot:GR3408265-4). Secondary antibodies included AF488 donkey anti-rabbit (1:1000, Jackson, Cat#: 711-545-152, Lot: 161527), AF647 donkey anti-rabbit (1:1000, Jackson, Cat#: 711-605-152, Lot:145270) AF568 donkey anti-rabbit (1:1000, Invitrogen, A10042, Lot: 2433862), AF488 donkey anti-goat (1:1000, Jackson, Cat#: 705-545-003, Lot: 146920) and AF568 donkey anti-goat (1:1000, Invitrogen, Cat#: A11057, Lot: 2160061).

### Alizarin Red S Staining

Brains after post-fixation of 4% PFA were embedded with paraffin, and cut into 5 μm-thick sections. After deparaffinization and rehydration, the tissue sections were stained with Alizarin Red S (G1038, Servicebio) for calcification labeling, and photographed with an Olympus BX43 light microscope. 40X objectives were utilized.

### Electron microscopy

Mice were intracardially perfused with 0.01 M PBS and 4% PFA and tissue with the size of 1 mm × 1 mm × 2 mm was collected from corpus callosum and then immersed in a mixture of 2.5% glutaraldehyde (GA) and 2% PFA for more than 2 hours at 4°C, washed two times in the phosphate buffer, then fixed with 1% osmium tetroxide. The specimen was dehydrated through a graded ethanol series, and then processed into araldite resin blocks. The specimen was placed in capsules contained embedding medium and heated at 60°C for 48hours. The samples were sectioned by a diamond blade (Leica EM UC7) to 70 – 90 nm thickness, followed by stained with uranyl acetate and lead citrate for contrast. All the specimens were viewed by transmission electron microscope (HITACHI H-7650).

### Microglia replacement by bone marrow transplantation (Mr BMT) in mice

For Mr BMT by busulfan, 8-week-old recipient mice were fed PLX5622 from day 0 to day 14. Then, the mice received busulfan (25 mg/kg of body weight each day) from day 9 to day 12 by intraperitoneal injection. Afterward, 1 × 10^7^ bone marrow cells harvested from the tibia and femur of the β-actin-GFP donor mouse were immediately introduced into the recipient mice on day 14 via intravenous injection. Then, the mice were fed a control diet. The mouse was fed neomycin (1.1 g/L) in acidic water (pH 2-3) throughout the procedure of microglia replacement.

### Traditional bone marrow transplantation (tBMT) in mice

Two approaches (irradiation-^13,20,21^ and busulfan-based^21^) were used to achieve myeloid ablation of tBMT in this study. For head-shielded tBMT by irradiation, 8-week-old recipient mice were fed CD from day 0 to day 14. Then, the heads of pretreated mice were shielded by lead and exposed to 9 Gy X-ray irradiation on day 14^13,20,21^. For tBMT by busulfan, 8-week-old recipient mice were fed CD from day 0 to day 14. Then, the mice received busulfan (25 mg/kg of body weight each day) from day 9 to day 12 by intraperitoneal injection. Afterward, 1 × 10^7^ bone marrow cells harvested from the tibia and femur of the β-actin-GFP donor mouse were immediately introduced into the recipient mice on day 14 via intravenous injection. Then, the mice were fed a control diet. The mouse was fed neomycin (1.1 g/L) in acidic water (pH 2-3) throughout the procedure of microglia replacement.

### Rotarod test

For the rotarod test, the mice were placed on an accelerating rotarod cylinder, the speed was slowly accelerated from 4 to 40 rpm within 5 min. and maintain for another 5 min at 40 rpm. A trial ended when the animal dropped or spun around for 3 consecutive revolutions without attempting to run on the rods. The speed of the rotarod when the mouse dropped was recorded. The test data are presented as the mean of speed (4 trials).

### Balance beam test

For the balance beam test, the mice were trained 2 days before the test. On day 3, the mice were placed on balance beam, and the number of slips and time of crossing the beam was measured. A trial was considered successful when the mice walked completely from one end of the beam to the other without pause. The test data are presented as the mean of a number of slips and time of crossing the beam (3 trials).

### Hindlimb clasping test

For the hindlimb clasping test, the mice were suspended by the tail and video-recorded for about 10 seconds. Hindlimb clasping was rated from 0 to 3 based on severity: be rated as 0 if hindlimbs splayed outward and away from the abdomen, be rated as 1 if one hindlimb retracted inwards towards the abdomen for at least 50% of the observation period, be rated as 2 if both hindlimbs partially retracted inwards towards the abdomen for at least 50% of the observation period, be rated as 3 if both hindlimbs completely retracted inwards towards the abdomen for at least 50% of the observation period.

### Single-cell RNA sequencing

The cell suspension was primarily according to our previous protocol^26^. In brief, the mice were perfused with 0.9% saline and inhibitor cocktail solution (actinomycin D 5 μg/mL, triptolide 10 μM, anisomycin 27.1 μg/mL), then the brains of mice were quickly sliced into 200 μm sections. Next, the sections were incubated in choline chloride solution with 10 μM NBQX (Tocris, Cat#: 0373), 50 μM APV (Tocris, Cat#: 3693), inhibitor cocktail, and influx with oxygen (95% O_2_ and CO_2_) for 30 min at 37 °C. After that, the sections were digested in 20 U/mL papain (Worthington, LK003178) and 100 U/mL DNase I (Sangon, B100649-0040) for 20 min at 37 °C, then the sections were incubated in 1mg/mL protease (Sigma-Aldrich, P5147) and 1 mg/mL dispase (Worthington, LS02106) for 20 min at 25 °C. Next, the sections were gently and repeatedly blown through pipette into single-cell suspensions, then filtered through 40 μm cell strainers. The cell debris was removed by 30% Percoll (Millipore, P1644) in DPBS with 0.5%BSA. Finally, the cells resuspended in DPBS with 0.5% BSA and approximately 24,000 cells for each group (N = 3 mice) were loaded for 10x Genomics library preparation. A cocktail of transcriptional and translational inhibitors was included in the cell preparation to prevent the *ex vivo* microglial reactivation^25,26^.

The single-cell libraries were constructed using the Chromium controller and Chromium single-cell 3 reagent version 2 kit (10x Genomics). Then the libraries were sequenced at the paired-end 150 bp using HiSeq X Ten (Illumina).

### Analysis of scRNA-seq data

Sample demultiplexing, barcode processing and single-cell 3’ gene counting were carried out with Cell Ranger 7.0.1 (10x Genomics). The 10-bp transcripts/UMI tags were extracted from the reads. Cellranger mkfastq with bcl2fastq 2.0.1 was applied to demultiplex raw base call files from the sequencer into sample-specific FASTQ files. Additionally, cellranger-compatible references were produced based on both genome sequences and transcriptome GTF files by using Cell Ranger mkref. These FASTQ files were aligned to the reference genome mm10 with cellranger count by using STAR 2.7.2. Aligned reads were then filtered for valid cell barcodes and UMI to generate filtered gene-barcode matrices.

Next, gene-barcode matrices were analyzed with Seurat 5.0.1. For quality control, low-quality cells (detected > 7,000 genes, < 200 genes and > 10% mitochondrial genes) were discarded. Filtered data were further normalized by SCTransform. For dimension reduction, data were reduction by using RunPCA with VariableFeatures, and the top 50 principal components (PCs) of the datasets were used for clustering and tSNE/UMAP visualization. Then the markers of each cluster were calculated by FindAllMarkers with resolution = 2.

To identify different brain cell types, we used multiple cell type-specific or enriched marker genes as previously described. In brief, these genes included but were not limited to *P2ry12*, *Tmem119*, *Hexb* and *Aif1* for microglia; *Ms4a7*, *Mrc1*, *Cx3cr1* and *Aif1* for macrophage; *Gfp* for Mr BMT derived cell; *Plp1*, *Mog*, *Mag* and *Mbp* for oligodendrocytes; *S100b*, *Gja1*, *Slc1a2* and *Aldh1l1* for astrocytes; *Syt1* and *Map2* for mature neurons, *Tubb3* and *Dcx* for immature neurons; *Syt1* and *Baiap3* for neuroendocrine cells (NendC); *Pdgfra* for OPCs; *Pdgfra*, *Dcn*, *Lama1* and *Lum* for vascular and leptomeningeal cells (VLMC); *Acta2* for vascular smooth muscle cells (VSMC); *Nkg7* and *Skap1* for nature killer cells; *Alas2* for hemoglobin-expressing vascular cells (Hb-VC); *Kcnj8* and *Vtn* for pericytes; *Cldn5* and *Pecam1* for endothelial cells (EC); *Npy* for olfactory ensheathing glia (OEG).

For DEG analysis, DEGs among groups of cells were identified using FindMarkers with parameters of min.pct = 0.1 and thresh.use = 0.25 (Wilcoxon rank-sum test). Genes with a |log2FC| > 0.25 and adjust P value < 0.05 were included and defined as DEGs. These DEGs were further used for GO and IPA analyses.

### GO analysis and IPA analysis

GO analysis was performed using the enrichGO function of the clusterProfiler 4.6.2 package, where categories with a false discovery rate (FDR) ≤ 0.05 were considered significantly enriched; categories were simplified using the ‘simplify’ function with a cutoff value = 0.7, and visualization was performed using the dotplot and barplot functions in clusterProfiler 4.6.2. DEGs were also subjected to IPA for functional analysis using the Ingenuity Knowledge Base (Qiagen Bioinformatics). The FC and FDR values of each gene were used to perform the core analysis, and pathways enriched with an adjusted P < 0.05 were considered significantly enriched.

### Statistics and reproducibility

The statistical approaches were indicated in figure legends. No statistical methods were used to pre-determine sample sizes but our sample sizes are similar to those reported in previous publications^13,21,26,40,41,47,64–66^. Mice were randomized for the naïve, Mr BMT and tBMT groups for experiment. Data distribution was assumed to be normal but this was not formally tested. Collection and analysis of scRNA-seq data were not performed blind to the conditions of the experiments. Collection of behavior experiment data was blind. Behavior data of mice with obesity, cataract or severe trauma were excluded from the analyses.

## Data availability

The data that support the findings of this study are available from the corresponding author Bo Peng at Fudan University within three months upon typical request.

## Acknowledgements

The authors thank Dr. Congcong Qi (Small Animal Behavior Monitoring Platform at the Laboratory Animal Resource Center, Fudan University) for the technical support in behavior tests and Jie Yang (Core Facility of Basic Medical Sciences at Shanghai Jiao Tong University School of Medicine) for the technical support in transmission electron microscopy. In addition, the authors express their gratitude and respect to all animals sacrificed in this study. This study was supported by STI2030-Major Projects (2022ZD0204700) (B.P.) and (2022ZD0207200 (Y.R.), National Natural Science Foundation of China (32170958) (B.P.), (82371255, 82071258) (L.C.) and 323B2030 (X.L.), Shanghai Pilot Program for Basic Research (Grant No. 21TQ014) (B.P.), “Shuguang Program” supported by Shanghai Education Development Foundation and Shanghai Municipal Education Commission (22SG07) (B.P.), Program of Shanghai Academic/Technology Research Leader (Grant No. 21XD1420400) (B.P.) and (23XD1402500) (L.C.), The Innovative Research Team of High-Level Local University in Shanghai (B.P.), Program for Outstanding Medical Academic Leader of Shanghai (2022LJ011) (L.C.), Shanghai Science and Technology Innovation Action Plan (23DZ2291500) (L.C.), Training Program for Research Physicians of Innovative Translational Ability supported by Shanghai Hospital Development Center (SHDC2022CRD037) (L.C.).

